# Folic acid-loaded chitosan nanoparticles: Cytotoxicity, acute and sub-acute oral toxicity and bioavailability in Wistar rats

**DOI:** 10.1101/2022.12.24.521870

**Authors:** Eram Fathima, Shashank Kumar, M M Patil, Farhath Khanum, T Anand

## Abstract

The study aims to develop folic acid-loaded chitosan nanoparticles (FA-Chi-NPs) as the vitamin is sensitive to heat and light and to investigate the safety of FA-Chi-NPs *in vitro* and *in vivo*. The prepared formulation showed an average diameter of 191.7 ± 2 nm with a zeta potential of + 52 ± 4 mV and PDI of 0.195 ± 0.03. The encapsulation efficiency was found to be 80%. *In vitro* cytotoxicity assays revealed that FA-Chi-NPs had no cytotoxic effect on Caco-2 cells. Acute-toxicity and subacute-toxicity of FA-Chi-NPs (0.4, 2, and 4 mg/kg body weight) revealed no mortality, and no gross morphological abnormalities were noticed. There were no dose-dependent toxic effects of free folic acid (FA) and FA-Chi-NPs in haematological, histological, plasma biochemical indices, oxidative stress enzymes, or other parameters when compared to the control group(s). The plasma and tissue levels of FA from FA-Chi-NPs were significantly higher. Results demonstrated enhanced bioavailability of FA from FA-Chi-NPs with no toxicity.

**Practical applications:** In the current study, nanoencapsulation is used as a potential approach to enhance folic acid bioavailability. The results showed that encapsulating folic acid in chitosan improved bioavailability with no apparent adverse effects, and thus chitosan could be a promising polymer for safe folic acid delivery.

## 1. INTRODUCTION

Encapsulation systems have been extensively explored for the oral delivery of a wide range of bioactive compounds, including various types of vitamins (Amiri et al., 2019; Azevedo et al., 2014; Chalella Mazzocato et al., 2019; de Melo et al., 2021; Estevinho et al., 2020; Fahami et al., 2018; Resende et al., 2020). Folic acid (FA; vitamin B_9_) is a vital micronutrient that plays a critical role in protein metabolism and nucleic acid biosynthesis, specifically during pregnancy and infancy, since it promotes rapid cell division and prevents neural tube abnormalities (Vora et al., 2021). FA has been encapsulated in a variety of delivery systems, including nano to microparticles, electrospun fibers, nano and microcapsules, hydrogels, and emulsions, for use in food formulations and nutraceuticals (Assadpour et al., 2016; Camacho et al., 2019; do Evangelho et al., 2019; Estevinho et al., 2020; Pérez-Masiá et al., 2015; Vora et al., 2020). FA is a synthetic form of folates that humans cannot produce and must therefore be obtained through diet (Pérez-Masiá et al., 2015). Furthermore, FA undergoes degradation reactions when exposed to temperature, light, acidic medium, and moisture (Ohrvik et al., 2011). As a result, encapsulating FA in delivery systems has resulted in a variety of important properties, including enhanced thermal and photostability slow and controlled delivery with improved bioavailability, making it more suitable for application in pharma and food formulations (Aceituno-Medina et al., 2015; Chapeau et al., 2017; Fu et al., 2018; Osojnik Črnivec et al., 2020; Penalva et al., 2015).

In this case, carbohydrate polymers have become beneficial as matrices for bioactive agent/drug carriers because of their biodegradability, biocompatibility, bioadhesibility, non-toxicity, higher relative abundance, and cost-effectiveness (Pateiro et al., 2021; Shukla et al., 2012). The use of natural polymer nanocarrier systems has demonstrated remarkable results in delivering hydrophilic or hydrophobic compounds with the desired shape, size, loading capacity, encapsulation efficiency, and controlled release to targeted tissue (Detsi et al., 2020). Among natural polymers, chitosan is widely used and an effective biocompatible nanocarrier with greater entrapment efficacies and controlled release characteristics, providing efficiency in the polymeric drug-delivery system (Govindappa et al., 2020). Chitosan interacts with sodium tripolyphosphate (STPP) through electrostatic forces because it is a cationic polysaccharide (Shu et al., 2002). This advantageous cross-linking mechanism not only aids in the encapsulation of biological/drug samples but also eliminates the need for toxic chemicals for cross-linking and emulsifying methods, which are typically hazardous to organisms (Berger et al., 2004).

In several earlier reports, FA has been used as a ligand to target cancer cells via conjugation with nanoparticle or surface coating. These nanoparticles were designed to improve tumor-targeted delivery of anti-cancer drugs like thymoquinone, doxorubicin hydrochloride, vincristine, protoporphyrin IX, docetaxel, and ligustrazine (Al-Nemrawi et al., 2022; Cheng et al., 2017; İnce et al., 2020; Salar et al., 2016; Song et al., 2013; Yang et al., 2010). A number of studies have been conducted to look into the effects of protein polymers (zein or casein) on FA bioavailability (Penalva et al., 2015; Peñalva et al., 2015). In our previous reports, optimization and characterization of FA-Chi-NPs were conducted, and the results revealed the stability of the prepared nanoformulation (Fathima et al., 2022). Moreover, to date, no report on the effect of chitosan polymer in improving FA bioavailability and safety aspects *in vivo* is available. Hence, the current research uses biodegradable chitosan polymer to develop FA-loaded chitosan nanoparticles (FA-Chi-NPs) since it is generally recognized as safe (GRAS) and biocompatible. However, FA-Chi-NPs may cause toxicity because they differ from bulk material in terms of high surface-to-volume ratio, nanosize, and ability to permeate effectively from the intestinal barrier to circulation. As a result, it is obligatory to investigate the safety aspects of FA-Chi-NPs developed in this work. However, there is research available on the physiological and pharmacological advantages of FA but minimal literature on the safety aspects of FA. No reports exist on the FA-loaded chitosan nanocarrier system. As a result, the present study focused on determining *in vitro* cytotoxicity using Caco-2 cell lines, the acute and subacute toxicity of FA-Chi-NPs, and the effect of prepared nanoparticles on hepatic and serum biochemical markers, haematological and histological changes, oxidative damage and bioavailability, and tissue distribution of FA from FA-Chi-NPs in Wistar rats. To substantiate the biochemical data, a histopathological examination was performed in the rat model. The findings reveal new evidence about the safe application of FA-Chi-NPs in the food and pharmaceuticals fields.

## 2. Materials and methods

### 2.1. Chemicals

Folic acid (vitamin B9, >99%), chitosan (Low molecular weight-80% deacetylated), Tween 80, Sodium tripolyphosphate (STPP), WST-1 (Water-soluble tetrazolium salt) and Dulbecco’s Modified Eagle’s Medium (DMEM), Trifluoroacetic acid (TFA), H_2_-DCFDA (2’,7’-Dichlorofluorescein diacetate), and Antibiotics were obtained from Sigma Aldrich (St. Louis, USA). Glacial acetic acid and Acetonitrile used are of HPLC grade procured from Merck Millipore (Millipore Sigma, USA). Caco-2 cell line was procured from NCCS, Pune, India. Fetal bovine serum (FBS) was purchased from Hi-media Laboratories (Mumbai, Maharashtra, India). All the analytical kits for urea, bilirubin, creatinine, albumin and total protein, alkaline phosphatase (ALP), Aspartate aminotransferase (AST), alanine aminotransferase (ALT), Lactate dehydrogenase (LDH), low-density lipoproteins (LDL), and high-density lipoproteins (HDL) were procured from Agappe Diagnostics (Ernakulam, India). ELISA kit for reduced glutathione (GSH), glutathione peroxidase (GPx), and glutathione reductase (GR) were purchased from Cayman Chemical Company (Ann Arbor, Michigan), ELISA kit for folic acid and lysis buffer were procured from Cloud Clone Corp (USA). All other solvents and reagents used were of analytical grade. All aqueous solutions used in this study were prepared using de-ionized Milli-Q water (Millipore, France).

### 2.2. Preparation of folic acid-loaded chitosan nanoparticles (FA-Chi-NPs)

The ionic gelation method previously reported by Alishahi et al. (2011) was employed to develop FA-Chi-NPs with marginal modification. The ionic interaction of positively charged chitosan and negatively charged STPP solution was used to prepare FA-Chi-NPs. Briefly, the protocol was standardized by dissolving low molecular weight chitosan (0.3%) in acetic acid (0.1%) under magnetic stirring overnight, resulting in a clear solution. FA (3 mg) and STPP (0.2%) were dissolved separately in milli-Q water, and the solutions were then filtered through 0.22 μm syringe filters. FA dissolved in 0.1 N NaOH solution and 0.5% Tween 80 were added dropwise to the chitosan solution and stirred for 30 min at 650 rpm at room temperature. The STPP was then added dropwise for another hour while magnetically stirring. A probe sonicator (PCI, India) was then used to sonicate the emulsion for 10 min at 60 Hz on ice. The prepared formulation was centrifuged at 12,000 rpm for 45 min at 4 °C and washed with milli-Q water several times. Meanwhile, the pellet was lyophilized and stored at 4 °C for further analysis.

### 2.3. Entrapment efficiency

The Encapsulation efficiency (EE) of FA was calculated by centrifuging the prepared nanoparticles at 12,000×g at 4 °C for 45 min (Sigma, Germany). The concentration of unencapsulated FA was analyzed by HPLC (Agilent Technologies Model 1260, USA) in comparison with standard FA by using the method outlined by Ilaiyaraja et al. (2020). The reverse-phase C-18 column (250 mm x 4.5 mm) was utilized for the analysis, and the mobile phase was 0.1% TFA solution and Acetonitrile with 0.1% TFA using a linear gradient system. The flow rate was set to 1 mL/min with the UV detector wavelength of 256 nm. The analysis temperature was kept at 30 °C. 20 μL of the sample was injected for analysis.

The entrapment efficiency (EE) was quantified by the equation given below:

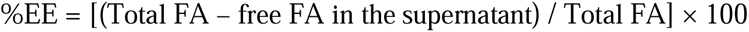

### 2.4. Particle size and zeta potential measurements

The particle size distribution and polydispersity index (PDI) of the obtained FA-Chi-NPs were measured using dynamic light scattering (DLS), and the zeta potential was determined using a Malvern Zetasizer (Nano ZS®, Malvern Instruments Ltd., UK). The prepared NPs were suspended in de-ionized water (1/4, v/v) to reduce the multiple scattering caused by the concentrated sample and analyzed under a 90° scattering angle at 25 °C in triplicate.

### 2.5. Evaluation of nanoparticles haemocompatibility

Hemolysis trials were carried out using a technique adapted from Jesus et al. (Jesus (2020). The human blood sample was obtained fresh from a healthy, non-smoking volunteer under institutional ethics protocols and explicit consent. In brief, 5 mL of blood was collected, the plasma was separated by centrifuging for 5 min at 2500 rpm, and the RBC (red blood cells) were isolated and repeatedly rinsed using normal saline. The extracted RBC was then reconstituted in normal saline to make a 25 mL suspension. To 2 mL of RBC suspension, 2 mL of different concentrations of FA-Chi-NPs suspended in normal saline were added (total concentrations were 200 µg/mL, 400 µg/mL, and 800 µg/mL). Negative (0% lysis) and positive (100% lysis) control samples were made by adding equal amounts of normal saline and 2% Triton X-100 to the RBC suspension, respectively. The samples were then incubated at 37 °C for 2, 4, and 6 h. The samples were gently shaken once every 30 min to resuspend the NPs and RBC. Following the incubation period, the samples were centrifuged for 5 min at 2500 rpm, and 1.5 mL of supernatant was collected and incubated at room temperature for another 30 min to facilitate haemoglobin oxidation. The absorbance of oxyhaemoglobin in supernatants was measured spectrophotometrically (Spectrostar Nano, BMG LABTECH) at 540 nm. The percentages of RBC hemolysis were calculated using the formula given below:

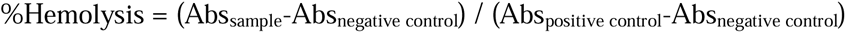

### 2.6. Evaluation of nanoparticles cytotoxicity

#### 2.6.1. Cell lines and culture conditions

Caco-2 (human intestinal adenocarcinoma cell line) was provided by the National Centre for Cell Sciences (NCCS), Pune, India, and was grown in DMEM media with supplements such as 10% fetal bovine serum, 2mM L-glutamine, and antimycotic solution (Sigma, USA) in a humidified incubator set at 37 °C and 5% CO_2_. The cells were grown in T-25 cm^3^ flasks under ideal conditions until they reached 60–70% confluency. Every other day, the culture media was changed, and the cells were allowed to differentiate for 6-7 days. Caco-2 cells at passages 10-20 were used.

#### 2.6.2. Water-soluble Tetrazolium Salt (WST-1) assay

Cytotoxicity was estimated by the WST-1, which analyzes the cell metabolic activity. Caco-2 cells were plated in 96 well plate at a density of 10^4^ cells/well and then incubated at 37 °C and 5% CO_2_ for 24 hr before treatment. The cells were then exposed to the different concentrations of free FA and FA-Chi-NPs and incubated for 24 hr at 37 °C. After the treatment, the plates were rinsed twice with PBS buffer and incubated with 10 µl of the WST-1 cell proliferation reagent along with 90 µl of fresh medium for 1 hr at 37 °C. The absorbance was measured at a 450 and 650 nm dual-wavelength using a Hidex plate chameleon TM V (Finland).

#### 2.6.3. Reactive oxygen species (ROS) generation

The levels of intracellular ROS in Caco-2 cells were determined using the method previously described by Chen et al. (2016), using H_2_-DCFDA (2’,7’-Dichlorofluorescein diacetate) with some modifications. In brief, 2 X10^4^ cells/well were seeded in a 96-well microplate and incubated for 24 hr in an incubator maintained at 37 °C and 5% CO_2_. The culture media was changed after the cells had reached 80% confluency, and the plate was incubated after treating the cells with various concentrations of free FA and FA-Chi-NPs (200 µg, 400 µg, and 800 µg). After 24 hr incubation, 100 µl of 20 µM H_2_-DCFDA was added and further incubated for 40 min at 37 °C. The positive control was prepared by treating control cells with a medium containing 200 mM H_2_O_2_ for 1 hr prior to the addition of H_2_-DCFDA. The medium containing H_2_-DCFDA was aspirated, and the cells were twice with PBS. After adding 100 µl of PBS, images were immediately captured by a fluorescence microscope (Olympus Corporation, Japan), and fluorescence intensity was read at λ_excitation_ and λ_emission_ of 485 nm and 535 nm, respectively, by Hidex plate chameleon TM V (Finland).

#### 2.6.4. Lipid peroxidation assay

Malondialdehyde (MDA), the lipid peroxidation product, was quantified according to Ohkawa et al. (1979). Briefly, Caco-2 cells were seeded in 6-well plates at a density of 5 − 10^5^ cells/well, followed by incubation at 37 °C. After reaching 80% confluence, the cells were treated with various concentrations of free FA and FA-Chi-NPs (200 µg, 400 µg, and 800 µg). After incubating 24 hr, the medium was removed, and the cells were collected, washed with PBS, and lysed with 1.15% KCl with 1% Triton X-100 by sonication. A 100 µL of the homogenate was mixed with 1.5 mL of 0.8 % thiobarbituric acid, 0.2 mL of 8.1% SDS, 1.5 mL of 20% acetic acid under acidic conditions (pH 3.5), and the final volume was made up to 4 mL by distilled water and boiled for 2 hr. Samples were cooled and centrifuged for 15 min at 2,000 rpm, the supernatants were collected, and the formation of TBARS was measured at 532 nm. Experiments were conducted in triplicates, and the results were expressed in nmol/mL of MDA.

### 2.7. Animals

Animal experiments were carried out in Wistar rats of both sexes after approval from the Institutional Animal Ethics Committee (IAEC-2019/NBT/35) of DRDO-Defence Food Research Laboratory, Mysore, India. Wistar rats (8 weeks old, 100 ± 12 g of body weight) were obtained from Vaarunya Biolabs Pvt. Ltd. (Bengaluru, Karnataka, India) and were fed with a standard pellet diet. Animals were housed individually in polypropylene cages in a temperature-controlled room (25 ± 2 °C) at a 12-hr light/dark cycle with < 60% relative humidity and provided ad libitum access to water and feed. Rats were maintained in an animal house facility for two weeks prior to the experiment and acclimatized to laboratory conditions.

#### 2.7.1. Acute oral toxicity

Rats were randomly divided into following 5 groups (n = 6, 3 male and 3 female) namely Group 1 – Control group (Physiological saline), Group 2 – (Free FA – 4 mg/kg body weight), Group 3 (FA-Chi-NPs – 0.4 mg/kg body weight), Group 4 (FA-Chi-NPs – 2 mg/kg body weight) and Group 5 (FA-Chi-NPs – 4 mg/kg body weight). Acute oral toxicity of FA-Chi-NPs was evaluated according to the Organization for Economic Co-operation and Development Guideline 420 (Acute toxic class method, OECD, 2001). After overnight fasting (∼12 hr), experimental rats were orally administered with a single dose of free FA (0.5 mL) at 4 mg/kg body weight, and FA-Chi-NPs (0.5 mL) at 0.4, 2, 4 mg/kg body weight, and the group received physiological saline was considered as control. On gavage, rats were observed for 14 days for any sign of mortality (changes in food consumption, body weight, clinical signs) and mortality (Mouhoub et al., 2018). At the end of the experiment, animals were anesthetized and sacrificed with an intraperitoneal injection of pentobarbital sodium (100 mg/kg body weight). Blood and organs were collected and studied for haematology, histology, biochemical parameters, and oxidative stress biomarkers.

#### 2.7.2. Sub-acute oral toxicity

Rats were randomly divided into 5 groups (n = 6, 3 male and 3 female) namely Group 1 – control group (physiological saline), Group 2 – (free FA – 4 mg/kg body weight), Group 3 – (FA-Chi-NPs – 0.4 mg/kg body weight), Group 4 (FA-Chi-NPs – 2 mg/kg body weight), and Group5 (FA-Chi-NPs – 4 mg/kg body weight). The sub-acute oral toxicity of free FA and FA-Chi-NPs was assessed according to the Organization for Economic Co-operation and Development Guideline 407 (Repeated dose 28-day oral toxicity study in rodents, OECD, 2008). Free FA and FA-Chi-NPs were orally gavaged once a day (0.5 mL) at 4 mg/kg body weight and 0.4, 2, 4 mg/kg body weight for 28 days, respectively, and the group with no treatment was taken as control. Throughout the experiment, rats were examined for any signs of mortality or morbidity. The groups which received free FA (Group 2) and FA-Chi-NPs (Group 3-5) were compared with the control group to observe a relationship with its plasma FA levels as a factor of bioavailability and toxicity. Animals were then sacrificed, and blood and tissues were collected for further investigation, as described above.

#### 2.7.3. Extraction of FA from plasma and tissues

Blood samples were collected from each animal separately and centrifuged at 3,000 rpm at 4 °C for 15 min, and plasma was collected into clean tubes and stored at −80 °C until further analysis. FA was extracted from tissues by homogenizing with a bead-mill homogenizer using lysis buffer (specific for ELISA, Cloud clone). Tissue homogenates were centrifuged for 15 min at 12,500 rpm, and the supernatant was separated and stored at −80 °C for further investigation. FA in plasma and tissues was estimated by the competitive inhibition ELISA method as per the manufacturer’s instruction (Cloud Clone Corp., USA).

#### 2.7.4. Food consumption, clinical signs, and body weight

Throughout the dosing, feed intake was documented daily, and body weight was recorded every week. Animals were also examined for any clinical signs of mortality and morbidity, such as variations in skin, fur, behaviour, nasal secretions, eyes, and mucous membranes. Besides these, lacrimation, piloerection, and abnormal respiratory patterns were all monitored daily.

#### 2.7.5. Haematological and Biochemical assays

At the end of each experiment, whole blood was collected in a tube containing ethylenediaminetetraacetic acid (EDTA) as an anticoagulant. Haematological parameters such as haemoglobin (HGB), white blood cells (WBC), lymphocyte, neutrophil, eosinophil, monocyte, basophil, packed cell volume (PVC), erythrocyte count (RBC), mean corpuscular volume (MCV), mean corpuscular haemoglobin (MCH), mean corpuscular haemoglobin concentration (MCHC), and platelets count were evaluated using the automated haematology analyzer (Medonic M32, Japan). The plasma glucose, triglycerides, cholesterol, total protein, albumin, globulin, urea, uric acid, creatinine, bilirubin, high-density lipoproteins (HDL), low-density lipoproteins (LDL), and enzyme activities such as aspartate aminotransferase (AST), alanine aminotransferase (ALT), alkaline phosphatase (ALP), lactate dehydrogenase (LDH) were investigated with standard assay kits.

#### 2.7.6. Histopathology and gross necropsy

After sacrificing the animals, organs such as brain, lungs, heart, liver, kidney, and intestine were harvested, and organ weights were recorded and processed for histological investigations. Histopathological examination was conducted using standard laboratory protocols. Organs were stored in 10% neutral formalin, and tissues were embedded in paraffin blocks. Sections were stained with hematoxylin and eosin (H and E) and analyzed microscopically for histological changes.

#### 2.7.7. Oxidative stress biomarkers

##### 2.7.7.1. Superoxide Scavenging activity

Superoxide dismutase (SOD) activity has been assessed by the modified protocol of Das et al. (2000). In brief, 3 mL of reaction mixture contained phosphate buffer (0.1 M pH-7.4), α-methionine (20 mM), EDTA (50 µM), hydroxylamine hydrochloride (10 mM), Triton-X 100 (1%), riboflavin (100 µM) was added to 100 µl of tissue homogenate. For control, a reaction mixture was used that contained 100 μl of distilled water instead of a sample, and the reaction mixture without riboflavin served as blank. The absorbance was recorded at a wavelength of 412 nm spectrophotometrically.

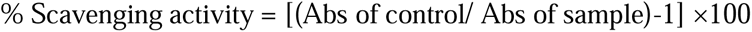

##### 2.7.7.2. Catalase activity

Catalase (CAT) activity was measured according to the method of Aebi (1984). In brief, by adding 50 µL of the homogenate to a 3 mL assay mixture comprising 8.8 mM H_2_O_2_ (1.5%) and 0.1 M sodium phosphate buffer of pH 7.4, the absorbance was measured at an interval of 20 sec for 3 min at 240 nm. The decrease in H_2_O_2_ level was expressed as U/min/mg of protein.

##### 2.7.7.3. Total antioxidant activity

The total antioxidant activity (TAC) was assessed using the phosphomolybdate method described by Prieto et al. (1999). Briefly, 1 mL of reagent solution containing 0.6 M sulphuric acid, 0.028 M sodium phosphate, and 0.004 M ammonium molybdate was added to 100 µL of tissue homogenate and incubated for 10 min at 95 °C. Upon cooling at room temperature, the absorbance was read at 695 nm against a blank, which contained a reagent solution with an appropriate volume of the solvent used for the sample. TAC was estimated using the ascorbic acid standard curve with a concentration of 0-200 μg/mL and expressed in μg of ascorbic acid equivalents per mg of protein.

##### 2.7.7.4. Lipid peroxidation

Lipid peroxidation (LP) was evaluated by the thiobarbituric acid reactive substances (TBARS) method (Ohkawa et al., 1979). In brief, 250 µL of the tissue homogenate was added to 1.5 mL of each 20% trichloroacetic acid and 0.6% thiobarbituric acid. The mixture was incubated in a boiling water bath for 30 min, cooled, and 2 mL of butanol was added, vortexed, and then centrifuged. The color of the butanol layer was measured at 535 nm spectrophotometrically and expressed in nM/mg of protein malonaldehyde (MDA).

##### 2.7.7.5. NO scavenging activity

Nitric oxide produced by sodium nitroprusside (SNP) solution reacts with oxygen at physiological pH to form nitrite ions, which can be measured using the Griess Illosyoy reaction (Green et al., 1982). Briefly, 1 mL of 5 mM SNP solution was mixed with 100 µL of the tissue homogenate and incubated for 60 min at 37 °C. Then 1.2 mL of Griess reagent (1% sulfanilamide in 5% H_3_PO_4_ and 0.1% α-naphthyl-ethylenediamine) was added. The absorbance of the pink chromophore was measured immediately at 550 nm. Ascorbic acid was taken as a control. Nitric oxide scavenging activity (%) was quantified using percent inhibition.

##### 2.7.7.6. GPx, GR, GSH activity

Glutathione peroxidase (GPx), glutathione reductase (GR), and reduced glutathione (GSH) activities were evaluated by assay kits from Cayman Chemical Company (Ann Arbor, Michigan) as per the manufacturer’s procedure. All enzymatic activities were normalized to the cellular protein content, as measured by a BCA protein assay kit (Sigma-Aldrich, USA).

### 2.8. Statistical analysis

All experiments were carried out in triplicate. The results were compared using a one-way analysis of variance (ANOVA) in SPSS software version 21.0, and Turkey’s test was used to determine whether the significant difference (p < 0.05) between the groups.

## 3. RESULTS

### 3.1. Characterization of FA-Chi-NPs

FA-Chi-NPs were successfully prepared using the ionic gelation method. DLS results illustrate the average particle size to be 191.6 ± 2 nm and zeta potential (ZP) to be + 51 ± 4 mV, demonstrating good stability and no aggregation NPs (Figure 1a). The polydispersity index (PDI) was found to be less than 0.2, indicating that the particles are of uniform size and shape. The entrapment efficiency (Figure 1b) of the NPs was found to be 80%.

**Figure 1.**
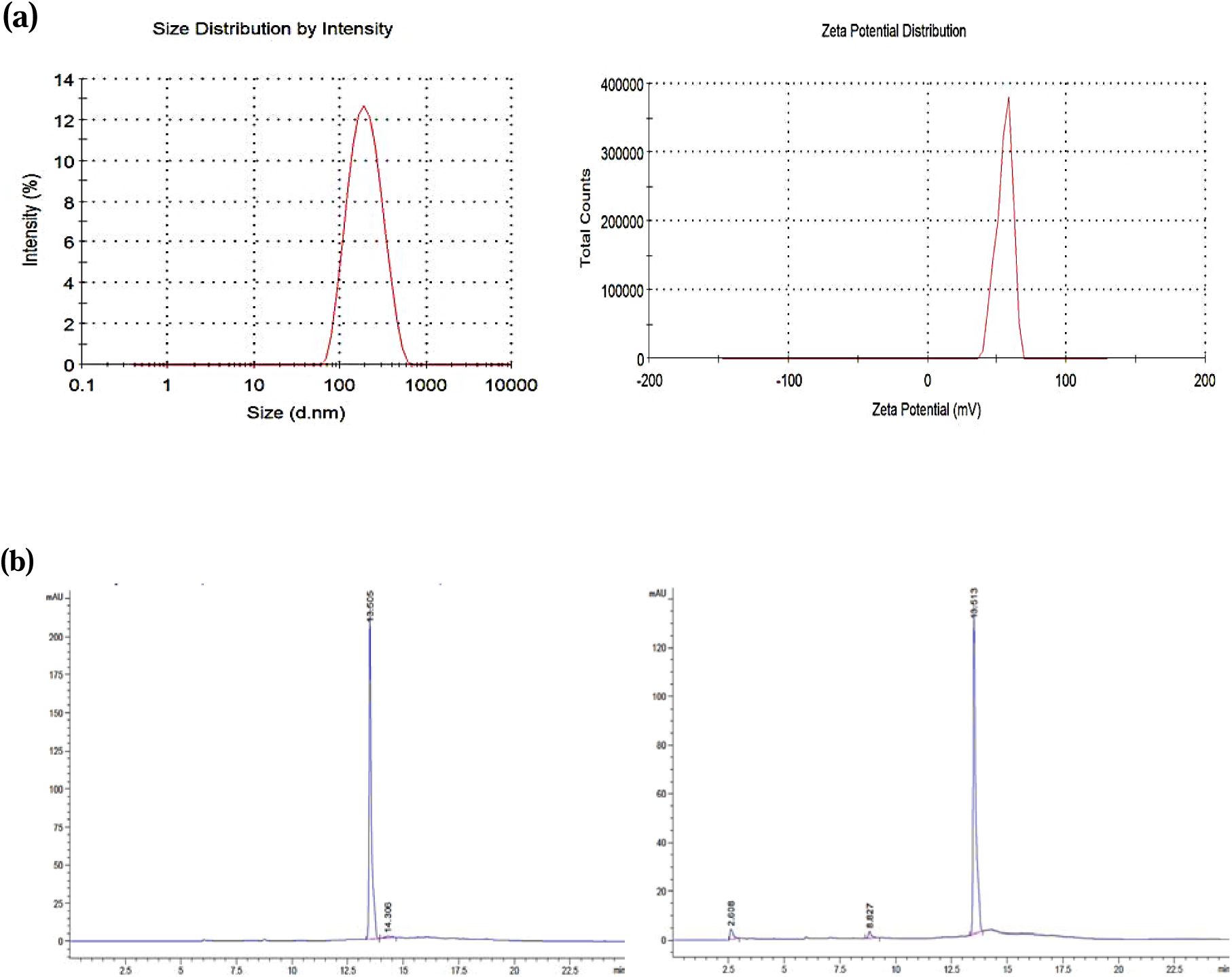
a) Size distribution graph; zeta potential graph of FA-Chi-NPs by zetasizer. b) HPLC chromatogram of FA standard and unencapsulated FA in the supernatant.

### 3.2. Haemocompatibility of FA-Chi-NPs

FA-Chi-NPs hemolytic activity was determined following 2, 4, and 6 h at 37 °C with RBCs. Results indicated that none of them generated more than 5% hemolysis, even at concentrations of 800 µg/mL. The percentage of hemolysis induced by nanoparticle treatment is represented in Figure 2.

**Figure 2.**
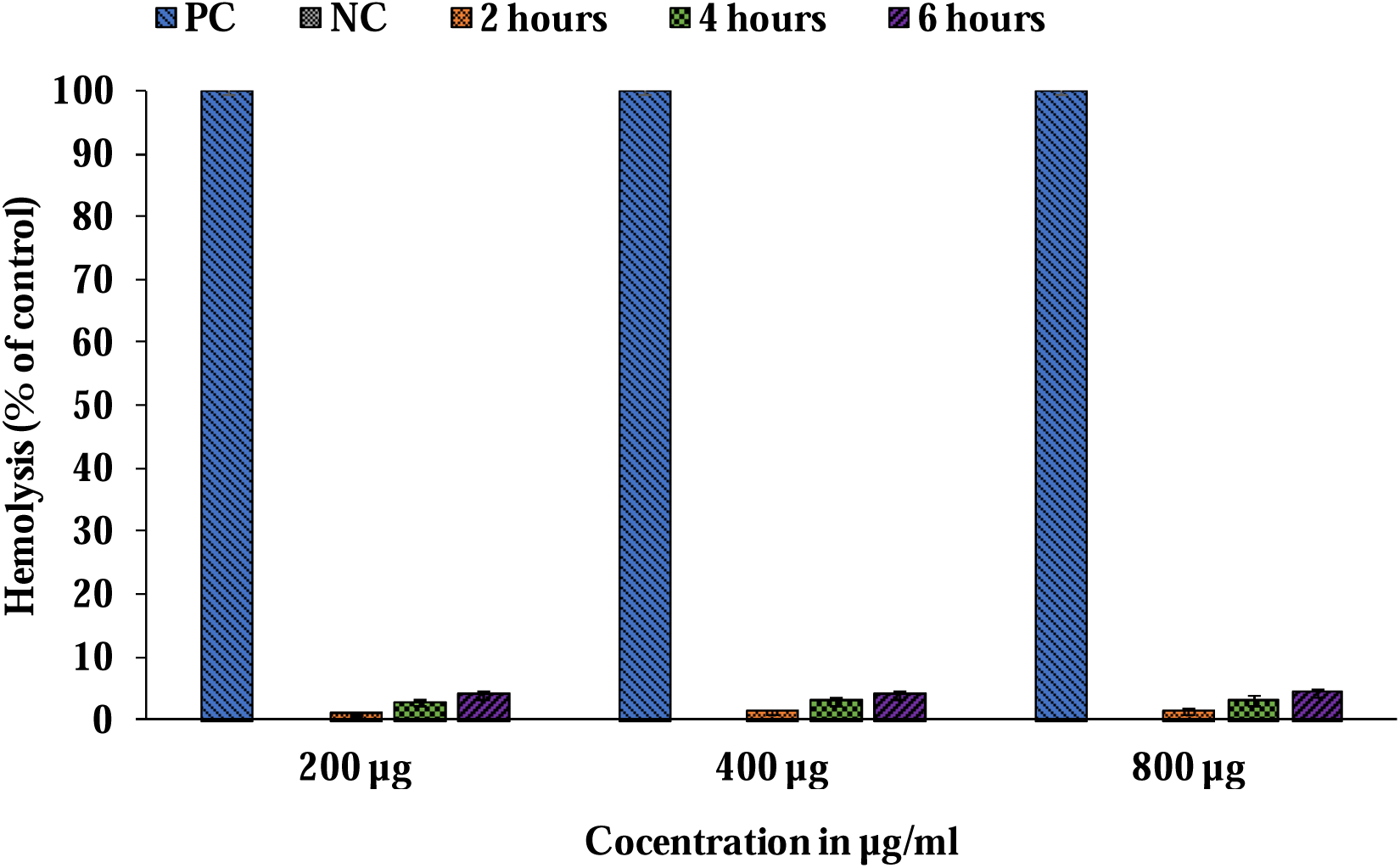
Hemolytic activity of FA-Chi-NPs in human blood after 2, 4, and 6 hr incubation at 37 °C. PBS and Triton-X-100 were used as negative control (NC) and positive control (PC), respectively. Results are expressed as mean ± SD, each performed in triplicates (n = 3).

### 3.3. *In vitro* cytotoxicity studies

#### 3.3.1. WST-1 assay

The cytotoxicity of synthesized FA-Chi-NPs was investigated by the WST-1 assay. We examined the cell viability of Caco-2 cells incubated with 10 different concentrations of free FA and FA-Chi-NPs (5, 10, 15, 25, 50, 100, 200, 400, and 800 μg/mL) for 48 hr (Figure 3a). Both free FA and FA-Chi-NPs do not display noticeable toxicity to Caco-2 cells as there was no significant cell death even after 48 hr of incubation at each of the tested concentrations.

**Figure 3.**
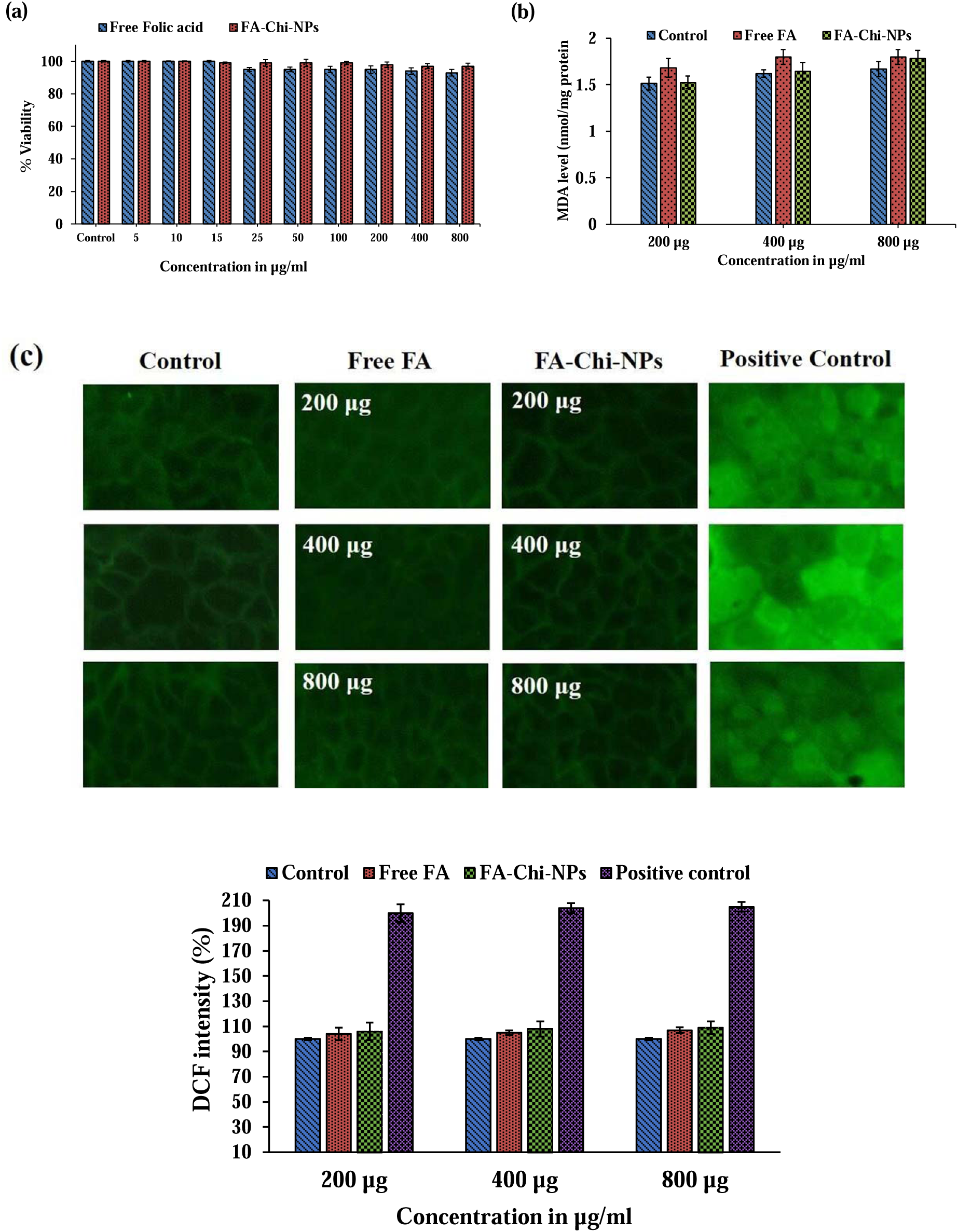
a) WST-1 assay on Caco-2 cell line treated with free FA and FA-Chi-NPs (n = 3) showing 95% cell viability up to the concentration ranging from 10 µg to 1000 µg/mL; b) Effect of free FA and FA-Chi-NPs on lipid peroxidation (MDA levels) in Caco-2 cell line; c) The effect of free FA and FA-Chi-NPs on reactive oxygen species (ROS) level in Caco-2 cell line are reported as DCF Fluorescence intensity (mean ± standard deviations). Positive control was prepared by treating normal cells with 200 mM H_2_O_2_ for 1 hr. Values are expressed in mean ± SE (n = 3).

#### 3.3.2. Intracellular reactive oxygen species (ROS) and lipid peroxidation

Reactive oxygen species (ROS) are unstable molecules, and their excessive production causes cell damage, which eventually leads to cell death (Schieber et al., 2014). Intracellular ROS production in Caco-2 cells in response to varied doses of free FA and FA-Chi-NPs was detected using 2’, 7’-Dichlorofluorescin diacetate (H_2_-DCFDA) assay (Figure 3c). This dye is non-fluorescent, but it turns fluorescent after cellular oxidation and acetate group removal by cellular esterases (Hanakova et al., 2014). Results revealed that FA-Chi-NPs did not increase ROS production in Caco-2 cells, demonstrating that the NPs did not induce oxidative stress (Figure 3c).

Lipid peroxidation was evaluated by estimating the formation of malondialdehyde (MDA) in the homogenates of Caco-2 cells. The exposure of cells to free FA and FA-Chi-NPs resulted in a non-significant increase in levels of MDA, and the results are summarized in Figure 3b.

### 3.4. Acute toxicity

Physiological and pathological signs of rats were studied during and after dosing with different concentrations of NPs. Acute oral doses of FA-Chi-NPs showed no evidence of mortality or toxicity associated with clinical signs and symptoms. The results showed that even at the highest dose of FA-Chi-NPs (4 mg/kg body weight) tested, there was no clinical evidence of toxicity associated with general health, behaviour, and locomotion of rats was observed either at the initial phase or throughout the 14-day post-treatment period, as compared with the control group. There were no significant changes (p < 0.05) in the water intake, daily feed, mean body weight, and organ weight (Figure 4 & Table 1) when compared to the control.

**Figure 4.**
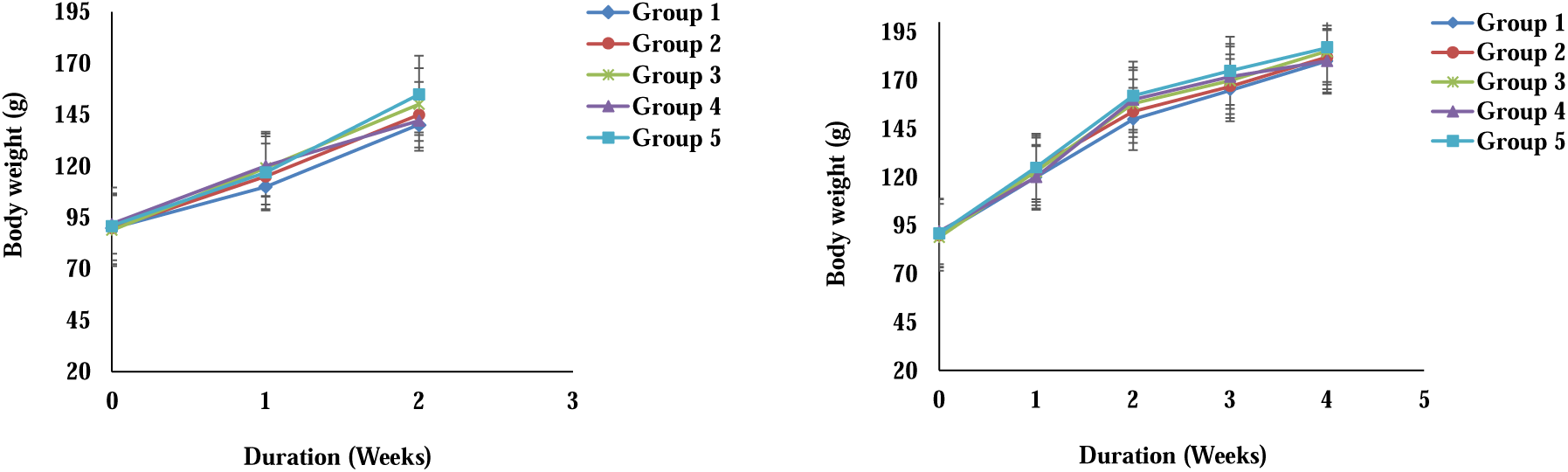
Body weight curves in rats gavaged with acute (a) and sub-acute (b) doses of FA-Chi-NPs. Group 1- control group (physiological saline), Group 2 (free FA - 4 mg/kg body weight), Group 3 (FA-Chi-NPs - 0.4 mg/kg body weight), Group 4 (FA-Chi-NPs - 2 mg/kg body weight), Group 5 (FA-Chi-NPs - 4 mg/kg body weight). Each value represents the mean ± SD (n = 6).

**Table 1.**
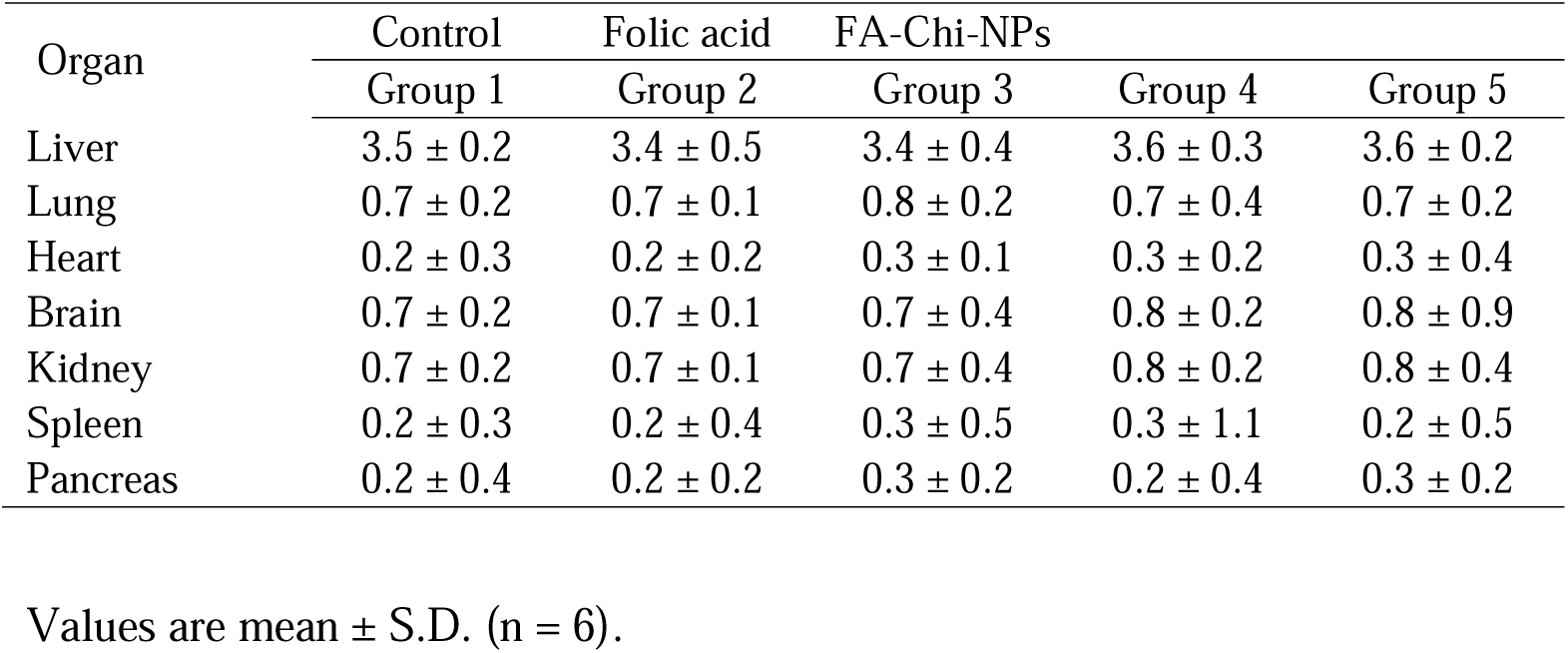
Relative organ weights (g/100 g) of rats administered with acute doses of FA and FA-Chi-NPs. Group 1- control group (physiological saline), Group 2 (free FA - 4 mg/kg mg/kg body weight), Group 3 (FA-Chi-NPs - 0.4 mg/kg body weight), Group 4 (FA-Chi-NPs - 2 mg/kg body weight), Group 5 (FA-Chi-NPs - 4 mg/kg body weight). Values are mean ± SD (n = 6).

Similarly, no treatment-related changes in haematological parameters were noticed in the treated group compared with the control group (Table 3). Various biochemical parameters were assessed using plasma to estimate the biochemical effect of NPs. The levels of ALT, AST, ALP, cholesterol, triglycerides, urea, creatinine, etc., were found normal, and no significant difference was detected in the plasma biochemical indices (Table 4). Ravi and Baskaran (2015) also reported that fucoxanthin-loaded chitosan nanogels did not cause any mortality or clinical signs of toxicity. All factors were found within the normal laboratory reference range for rats, with no variations among the experimental groups.

**Table 2.**
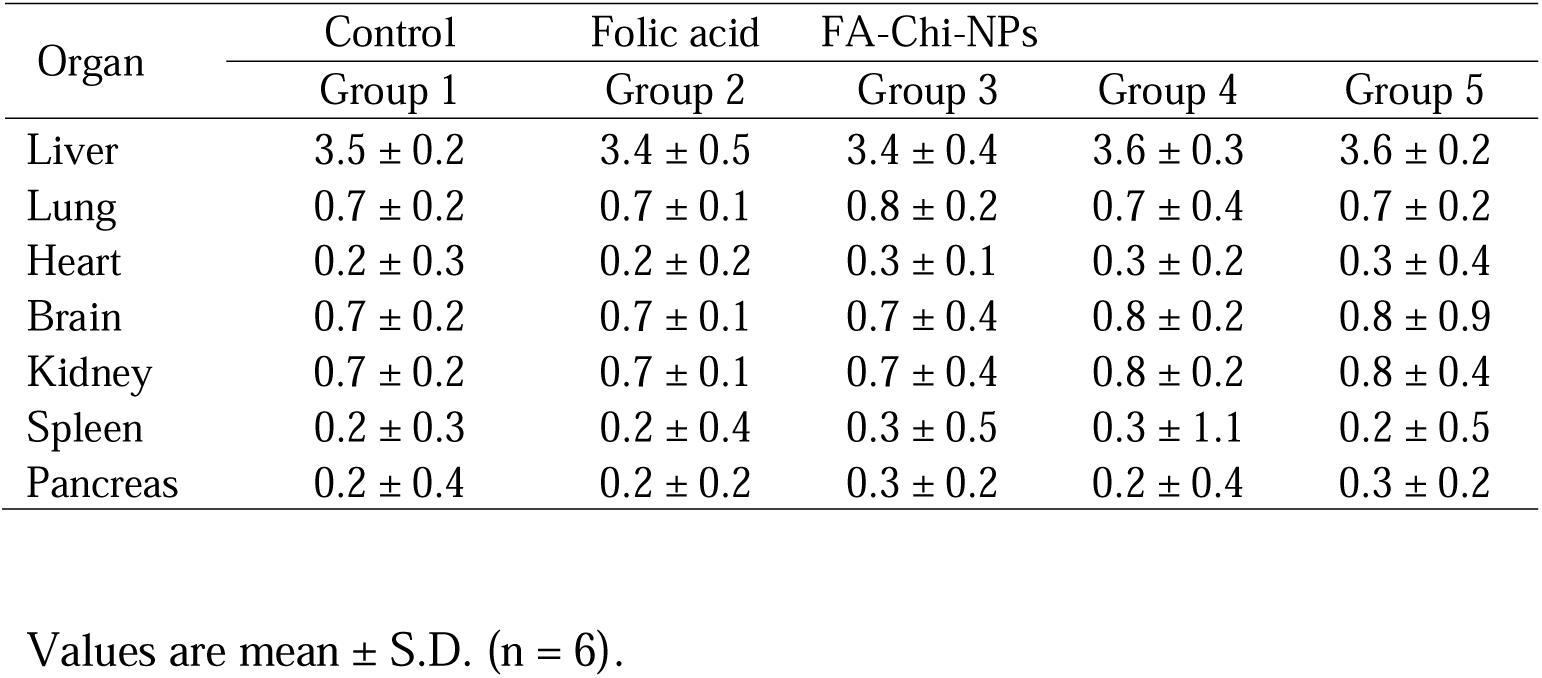
Relative organ weights (g/100 g) of rats administered with sub-acute doses of FA and FA-Chi-NPs Group 1-control group (physiological saline), Group 2 (free FA - 4 mg/kg body weight), Group 3 (FA-Chi-NPs - 0.4 mg/kg body weight), Group 4 (FA-Chi-NPs - 2 mg/kg body weight), Group 5 (FA-Chi-NPs - 4 mg/kg body weight). Values are mean ± SD (n = 6).

**Table 3.**
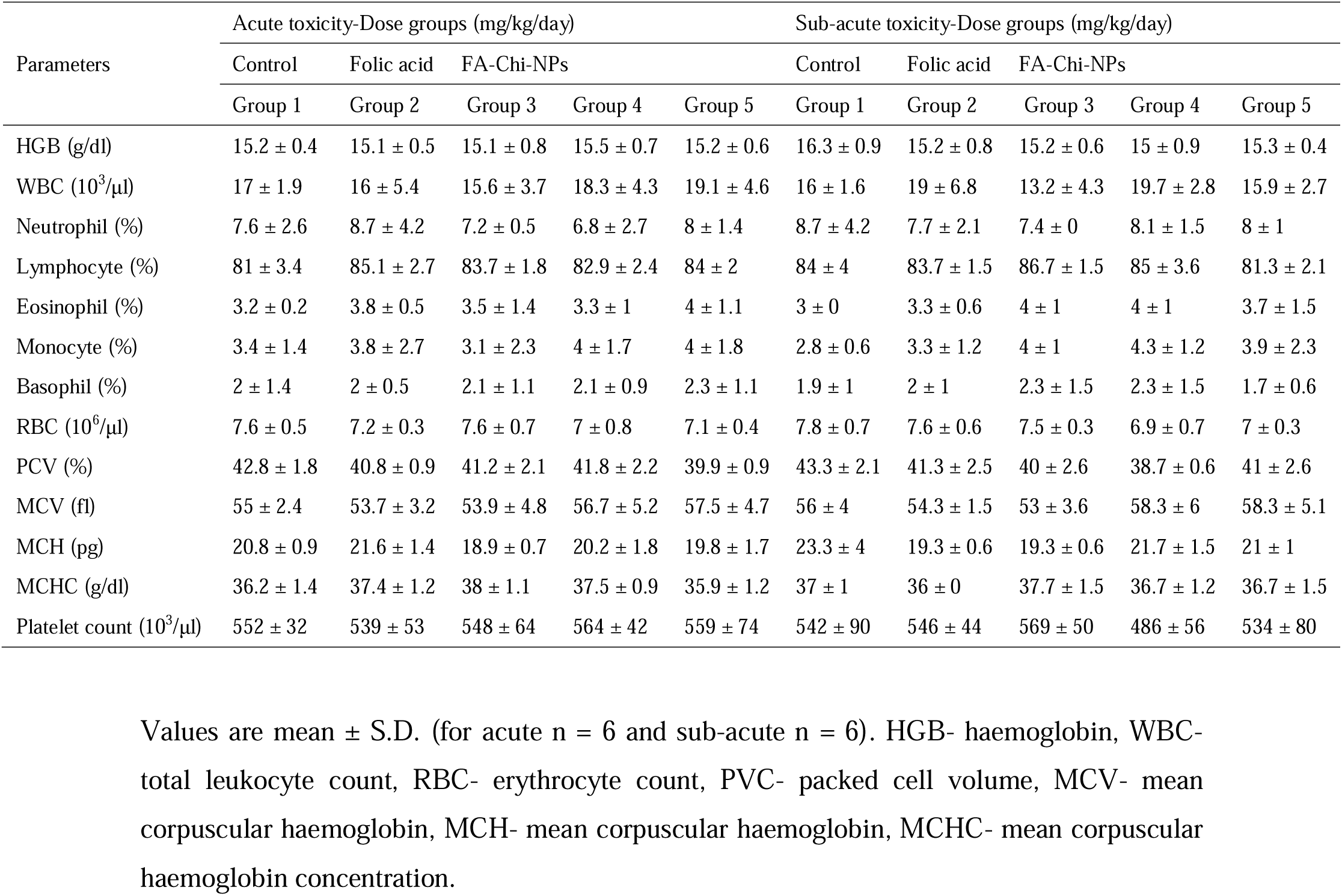
Haematological parameters of rats administered with acute and sub-acute doses of FA and FA-chi-NPs. Group 1-control group (physiological saline), Group 2 (free FA - 4 mg/kg body weight), Group 3 (FA-Chi-NPs - 0.4 mg/kg body weight), Group 4 (FA-Chi-NPs - 2 mg/kg body weight), Group 5 (FA-Chi-NPs - 4 mg/kg body weight). Values are mean ± S.D. (for acute n = 6 and sub-acute n = 6). HGB-haemoglobin, WBC- total leukocyte count, RBC- erythrocyte count, PVC- packed cell volume, MCV- mean corpuscular haemoglobin, MCH- mean corpuscular haemoglobin, MCHC- mean corpuscular haemoglobin concentration.

**Table 4.** Effect of acute and sub-acute dose of FA and FA-Chi-NPs on plasma biomarkers reflecting liver, kidney, and heart functions of rats. Group 1- control group (physiological saline), Group 2 (free FA - 4 mg/kg body weight), Group 3 (FA-Chi-NPs - 0.4 mg/kg body weight), Group 4 (FA-Chi-NPs - 2 mg/kg body weight), Group 5 (FA-Chi-NPs - 4 mg/kg body weight). Values are mean ± SD (for acute n = 6 and sub-acute n = 6). AST- aspartate aminotransferase, ALT- alanine aminotransferase, ALP- alkaline phosphatase, LDL- low density lipoproteins, HDL- high density lipoproteins, LDH-lactate dehydrogenase (p < 0.05).

| Parameters | Acute toxicity-Dose groups (mg/kg/day) |  |  |  |  | Sub-acute toxicity-Dose groups (mg/kg/day) |  |  |  |  |
| --- | --- | --- | --- | --- | --- | --- | --- | --- | --- | --- |
|  | Control | Folic acid | FA-Chi-NPs |  |  | Control | Folic acid | FA-Chi-NPs |  |  |
|  | Group 1 | Group 2 | Group 3 | Group 4 | Group 5 | Group 1 | Group 2 | Group 3 | Group 4 | Group 5 |
| Cholesterol (mg/dL) | 43.7 ± 5.7 | 45.8 ± 3.5 | 45.5 ± 7.2 | 44.3 ± 3.7 | 43.9 ± 2.7 | 44.2 ± 5.2 | 46.5 ± 7.1 | 43.8 ± 2.7 | 45.3 ± 7.4 | 44.5 ± 5.4 |
| Triglycerides (mg/dL) | 80.2 ± 3.8 | 81.2 ± 9.2 | 86.3 ± 7.4 | 75.5 ± 4.3 | 81.3 ± 8.3 | 90.5 ± 5.4 | 98.8 ± 7.8 | 86.3 ± 5.1 | 89.5 ± 3.5 | 91.3 ± 6.8 |
| Glucose (mg/dL) | 98.3 ± 2.7 | 100.2 ± 5.1 | 99.5 ± 5.1 | 105 ± 5.9 | 99.8 ± 7.2 | 103 ± 4.1 | 100.2 ± 2.8 | 105.1 ± 1.8 | 100.5 ± 6.7 | 101.2 ± 2.5 |
| AST (U/L) | 28.2 ± 2.8 | 30.7 ± 6.4 | 29.6 ± 2.3 | 31.5 ± 1.9 | 30.8 ± 2.8 | 29.2 ± 1.4 | 30.4 ± 3.2 | 32.1 ± 1.5 | 30.7 ± 4.1 | 28.8 ± 3.7 |
| ALT (U/L) | 22.5 ± 5.4 | 20.5 ± 1.9 | 21.5 ± 2.8 | 22.3 ± 4.1 | 20.5 ± 2.7 | 24.4 ± 4.2 | 25.3 ± 6.5 | 23.8 ± 4.6 | 22.5 ± 3.8 | 23.6 ± 4.3 |
| Albumin (g/dL) | 2.9 ± 0.2 | 3 ± 0.1 | 2.7 ± 0.2 | 2.8 ± 0.2 | 3.1 ± 0.4 | 3.2 ± 0.1 | 3.5 ± 0.1 | 3.6 ± 0.3 | 3.4 ± 0.8 | 3.2 ± 0.3 |
| Total protein (g/dL) | 5.2 ± 0.3 | 5.5 ± 0.7 | 5.1 ± 0.2 | 5.6 ± 0.4 | 5.1 ± 0.8 | 6.5 ± 0.8 | 6.1 ± 0.2 | 6.8 ± 0.6 | 6.1 ± 0.1 | 6.9 ± 0.4 |
| ALP activity (U/L) | 220 ± 10 | 232 ± 8.5 | 212 ± 15.2 | 240 ± 7.3 | 235 ± 14.3 | 250 ± 9.4 | 265 ± 12.6 | 245 ± 10.2 | 260 ± 7.5 | 245 ± 9.8 |
| Total Bilirubin (mg/dL) | 0.5 ± 0.1 | 0.52 ± 0.3 | 0.6 ± 0.4 | 0.58 ± 0.1 | 0.52 ± 0.1 | 0.57 ± 0.3 | 0.54 ± 0.2 | 0.61 ± 0.1 | 0.61 ± 0.5 | 0.59 ± 0.7 |
| Urea (mg/dL) | 28.79 ± 6.9 | 26.63 ± 5.3 | 25.62 ± 1.2 | 28.12 ± 3.4 | 25.96 ± 7.7 | 46.96 ± 2.8 | 45.56 ± 4.3 | 47.88 ± 5.2 | 48.82 ± 4.7 | 46.14 ± 2.6 |
| Creatinine (mg/dL) | 0.5 ± 0.1 | 0.4 ± 0.1 | 0.4 ± 0.2 | 0.5 ± 0.1 | 0.5 ± 0.1 | 1.1 ± 0.2 | 1.3 ± 0.1 | 1.4 ± 0.3 | 1.2 ± 0.1 | 1.3 ± 0.3 |
| Uric acid (mg/dL) | 0.35 ± 0.1 | 0.32 ± 0.1 | 0.31 ± 0.2 | 0.32 ± 0.2 | 0.34 ± 0.1 | 0.31 ± 0.2 | 0.28 ± 0.1 | 0.27 ± 0.3 | 0.32 ± 0.2 | 0.29 ± 0.2 |
| HDL-C (mg/dL) | 51 ± 7.3 | 54 ± 8.2 | 53 ± 11.2 | 57 ± 15.6 | 52 ± 20 | 71 ± 10 | 74 ± 13 | 78 ± 0.9 | 75 ± 8 | 70 ± 14 |
| LDL-C (mg/dL) | 45 ± 0.8 | 47 ± 0.6 | 44 ± 1.2 | 47 ± 2.6 | 48 ± 12 | 75 ± 6.5 | 71 ± 3.8 | 69 ± 14 | 78 ± 9.8 | 77 ± 6.4 |
| LDH (U/L) | 110 ± 15.6 | 108 ± 21 | 105 ± 23.2 | 108 ± 19 | 105 ± 10.2 | 98 ± 15.9 | 95 ± 21.5 | 97 ± 31.1 | 98 ± 20.6 | 96 ± 18.6 |

Furthermore, histopathology of the brain, lungs, heart, liver, kidney and intestine of FA-Chi-NPs fed rats was also normal compared with control (Figures 5a-f). Hence, administration of FA-Chi-NPs up to 4 mg/kg body weight has no adverse implications in rats.

**Figure 5.**
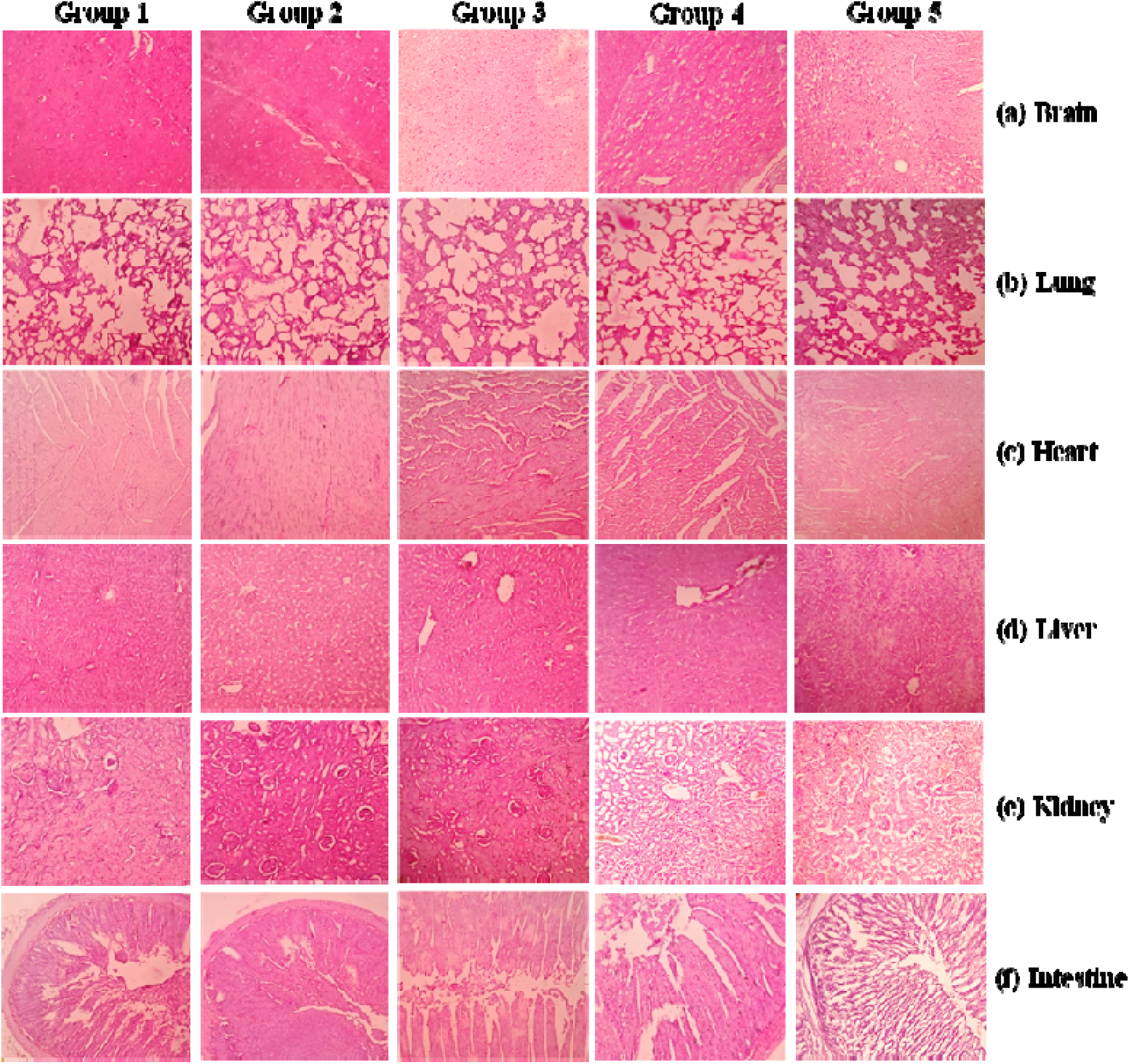
Histology of the brain, lungs, heart, liver, kidney, and intestine after an acute dose of free FA and FA-Chi-NPs for 14 days. Group 1- control group (physiological saline), Group 2 (free FA - 4 mg/kg body weight), Group 3 (FA-Chi-NPs - 0.4 mg/kg body weight), Group 4 (FA-Chi-NPs - 2 mg/kg body weight), Group 5 (FA-Chi-NPs - 4 mg/kg body weight). The observation was made at 40x magnification, H and E staining.

### 3.5. Sub-acute toxicity

#### 3.5.1. Clinical observations

During the sub-acute toxicity assessment, no treatment-related pathological signs or mortality were observed in any free FA, or FA-Chi-NPs fed group throughout the 28-day study. The average food intake per day was 8-12 g. In comparison to the control group, there were no dose-dependent changes in the body weight of experimental rats (Figure 4b). Additionally, abnormalities such as constipation, diarrhea, or other gastrointestinal disorders were not detected in any of the tested groups. A necroscopy investigation confirmed no macroscopic change in the heart, thoracic, or abdominal cavities’ outer surface or other orifices in both males and females.

#### 3.5.2. Haematology studies

Haematological parameters are used to detect physiological and pathological alterations in animals and humans. The sub-acute effect of FA-Chi-NPs on haematological indices is depicted in Table 3. A non-significant alteration was evidenced in HGB, RBC, WBC, MCV, MCHC, platelet count, and polymorphs, including basophils, eosinophils, neutrophils, monocytes, and lymphocytes in treated rats compared with the control. According to the current findings, repeated dosages of FA-Chi-NPs up to 4 mg/kg body weight for 28 days had no toxic effect on the haematological parameters tested. These results indicate that the nanoparticle formulations are safe at 4 mg/kg body weight.

#### 3.5.3. Plasma chemistry

The plasma biochemical indices of the present study exhibiting liver, kidney, and heart functioning are summarized in Table 4. Administration of FA-Chi-NPs (0.4, 2, 4 mg/kg body weight) had no significant difference between the control groups and experimental groups. Liver enzymes such as AST ALP or ALT levels were not changed in a dose-dependent manner (p < 0.05) in NPs fed group. Moreover, the alteration (12%) in the AST activity in the chitosan-fucoxanthin nanogels treatment is reported by Ravi and Baskaran (2015) but was not validated by histology of the liver or specific liver functional indicators in plasma such as ALP and bilirubin. However, the alterations were within the normal physiological range of AST and thus not related to nanogels. No significant alteration in the renal functional markers such as uric acid, urea, or creatinine levels, demonstrating no changes in the kidney function of male and female rats. The findings of plasma chemistry examination from treatment groups show that administration of the FA-Chi-NPs at doses up to 4 mg/kg/day to rats for 28 days had no toxicologically significant harmful effects.

#### 3.5.4. Gross necroscopy, organ weight, and histopathology

The macroscopic evaluation demonstrated no sign of abnormalities, necroscopy, or lesions in the FA-Chi-NPs treated groups compared to the control group. Additionally, there were no treatment-related changes in the relative organ weight of FA-Chi-NPs administered groups of acute and sub-acute doses, indicating no adverse impacts (Table 1 and Table 2). Histopathological tests were conducted to determine the influence of FA-Chi-NPs on vital organs such as the brain, lungs, heart, liver, kidney, and intestine (Figures 6a-f).

**Figure 6.**
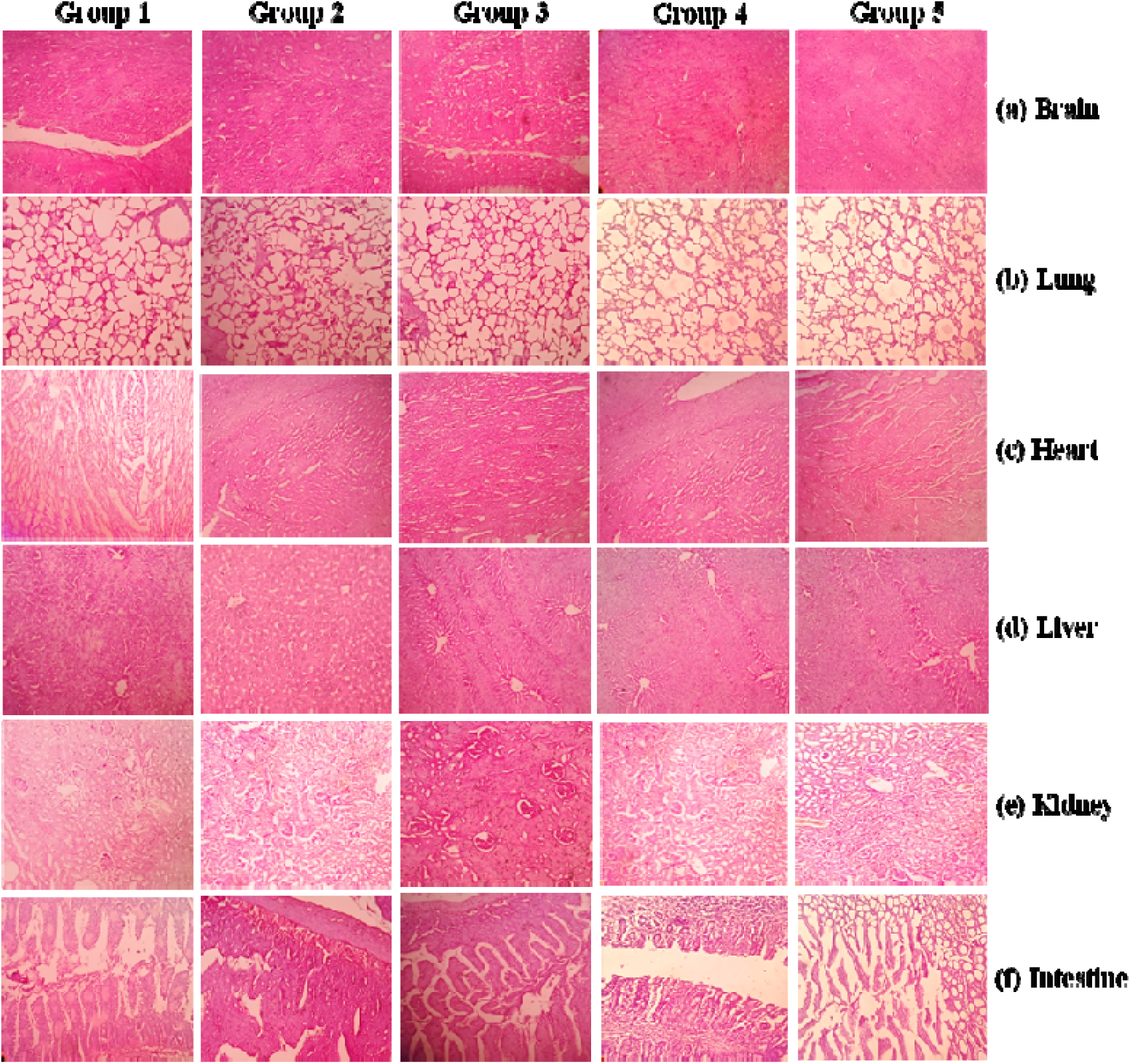
Histology of the brain, lungs, heart, liver, kidney, and intestine after a sub-acute dose of free FA and FA-Chi-NPs for 28 days. Group 1- control group (physiological saline), Group 2 (free FA - 4 mg/kg body weight), Group 3 (FA-Chi-NPs - 0.4 mg/kg body weight), Group 4 (FA-Chi-NPs - 2 mg/kg body weight), Group 5 (FA-Chi-NPs - 4 mg/kg body weight). The observation was made at 40x magnification, H and E staining.

Brain sections studied showed normal brain tissue with foci of neuronal fibres and blood vessels. Occasional lymphocytes were seen with foci of a normal axonal and dendritic histologic pattern. No neuronal degeneration was observed (Figure 6a). Lung Sections investigated showed normal alveoli with intervening alveolar blood vessels and bronchiolar features. The alveolar wall displayed normal orientation and interalveolar features. There was no evidence of alveolar wall deformation or congestion (Figure 6b). Sections of the heart studied exhibited normal cardiac muscle fibers in a transversely oriented pattern with foci of muscle cells with normal orientation and polarity. Hence, no cellular degeneration/distortion was seen (Figure 6c). Hepatic tissue histology is normal in the examined liver sections. Hepatocyte foci in hexagonal lobules with central vein, peripheral portal triad, foci in cords with normal sinusoidal space, and minor lymphocytic infiltration were seen. There was no evidence of hepatocytic degeneration (Figure 6d). Kidney Sections studied show predominantly normal histology of the kidney and foci of glomeruli and tubules with normal differentiation and polarity. No glomerular capillary distortion was seen (Figure 6e). Sections of the intestine studied show normal intestinal histology with intervening benign glandular villi. Foci of columnar cells with evenly distributed goblet cell lining the villi were observed. Thus, Individual cells show normal polarity and orientation (Figure 6f). As a result, histopathological examinations were found to have no pathologies or signs of organ damage even at higher concentrations than the control group.

#### 3.5.5. Oxidative stress biomarkers

*In vivo* nanotoxicological investigations using various routes of administration have a higher impact on determining the adverse biological effects (Reddy et al., 2017). Thus, the impact of free FA and FA-Chi-NPs on the antioxidant parameters was studied in the liver, kidney, and spleen of experimental animals for both acute (Figures 7a-h) and sub-acute doses (Figures 8a-h). In the current investigation, all of the treated groups showed non-significant increases in MDA, NO, GSH, TAC, SOD, CAT, GPx, and GR activity.

**Figure 7.**
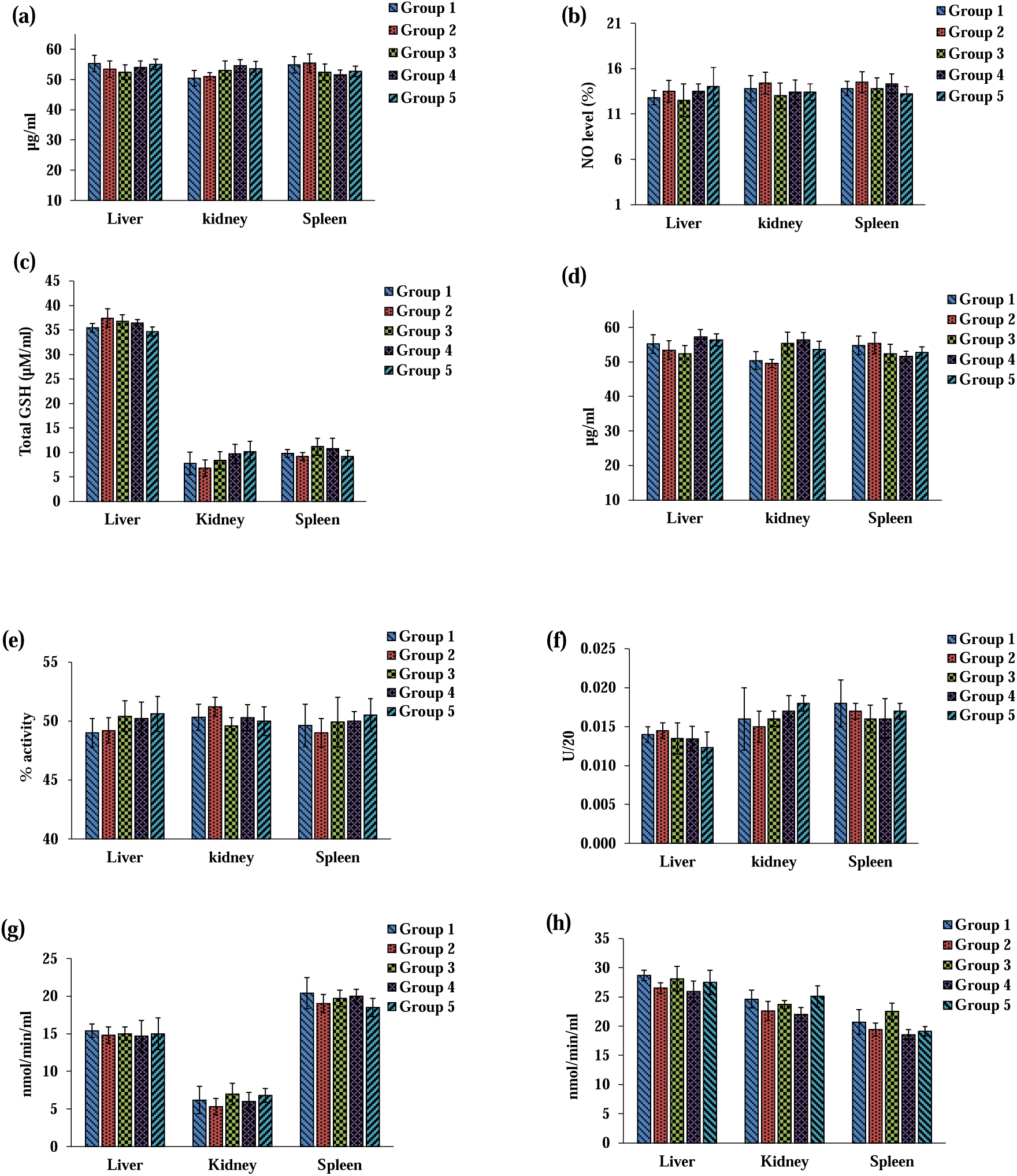
Indicators of oxidative stress in rat liver, kidney, and spleen tissues following acute dose administration of free FA and FA-Chi-NPs. Group 1-control group (physiological saline), Group 2 (free FA - 4 mg/kg body weight), Group 3 (FA-Chi-NPs - 0.4 mg/kg body weight), Group 4 (FA-Chi-NPs - 2 mg/kg body weight), Group 5 (FA-Chi-NPs - 4 mg/kg body weight). Levels of (a) LP, (b) NO, (c) GHS, (d) TAC, (e) SOD, (f) CAT, (g) GPx, and (h) GR were measured in the control and FA-Chi-NPs administered groups. Data are expressed as mean ± SD (n = 6). Statistical significance (p < 0.05) was determined by one-way analysis of variance (ANOVA) and multiple comparisons conducted using Tukey’s test.

**Figure 8.**
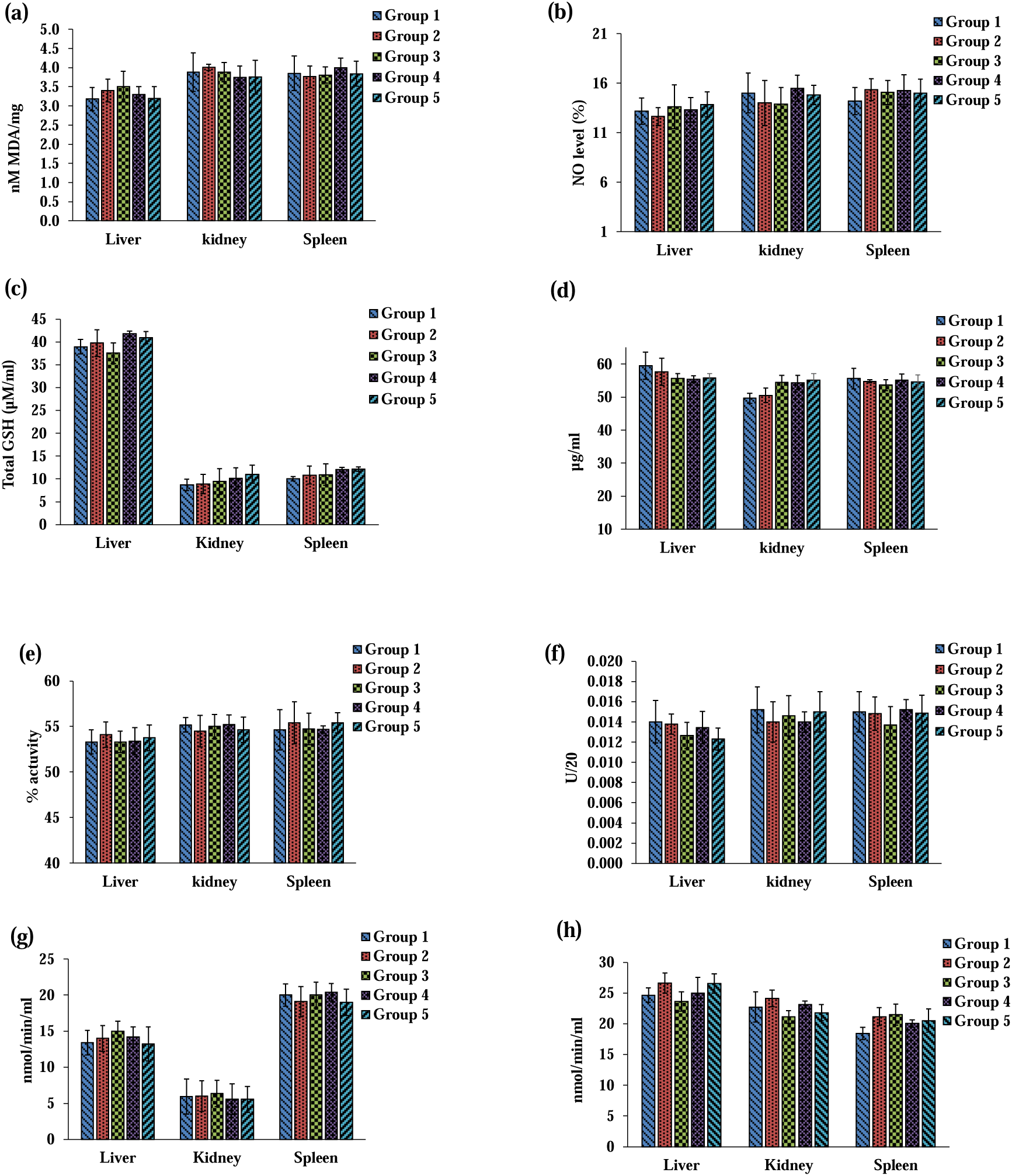
Indicators of oxidative stress in rat liver, kidney, and spleen tissues following sub-acute dose administration of free FA and FA-Chi-NPs. Group 1-control group (physiological saline), Group 2 (free FA - mg/kg body weight), Group 3 (FA-Chi-NPs - 0.4 mg/kg body weight), Group 4 (FA-Chi-NPs - 2 mg/kg body weight), Group 5 (FA-Chi-NPs - 4 mg/kg body weight). Levels of (a) LP, (b) NO, (c) GHS, (d) TAC, (e) SOD, (f) CAT, (g) GPx, and (h) GR were measured in the control and FA-Chi-NPs administered groups. Data are expressed as mean ± standard deviation (n = 6). Statistical significance (p < 0.05) was determined by one-way analysis of variance (ANOVA) and multiple comparisons conducted using Tukey’s test.

#### 3.5.6. FA bioavailability and tissue distribution

The plasma concentration profiles and tissue distribution of the FA after oral gavage (sub-acute doses) of free FA and FA-Chi-NPs are depicted in Figures 9a & 9b. Sub-acute doses of free FA (4 mg/kg body weight) to rats showed increased FA levels up to 85 ng/mL in the plasma and FA-Chi-NPs (0.4, 2, 4 mg/kg body weight) also displayed a dose-dependent elevation of FA in the plasma (p < 0.05) 60.1, 180, 300.2 ng/mL compared to the control group. The tissue distribution study of FA from FA-chi-NPs is represented in Figure 9b. The highest deposition of FA in the group fed with free FA (4 mg/kg body weight) and FA-Chi-NPs (0.4, 2, 4 mg/kg body weight) was detected in the jejunum (160 and 36, 200, 381 ng/g), duodenum (310 and 66, 370, 650 ng/g), and liver (112 and 28, 127, 268 ng/g) in comparison with the control group. The lowest quantities of FA were detected in heart, lung, and spleen tissues.

**Figure 9.**
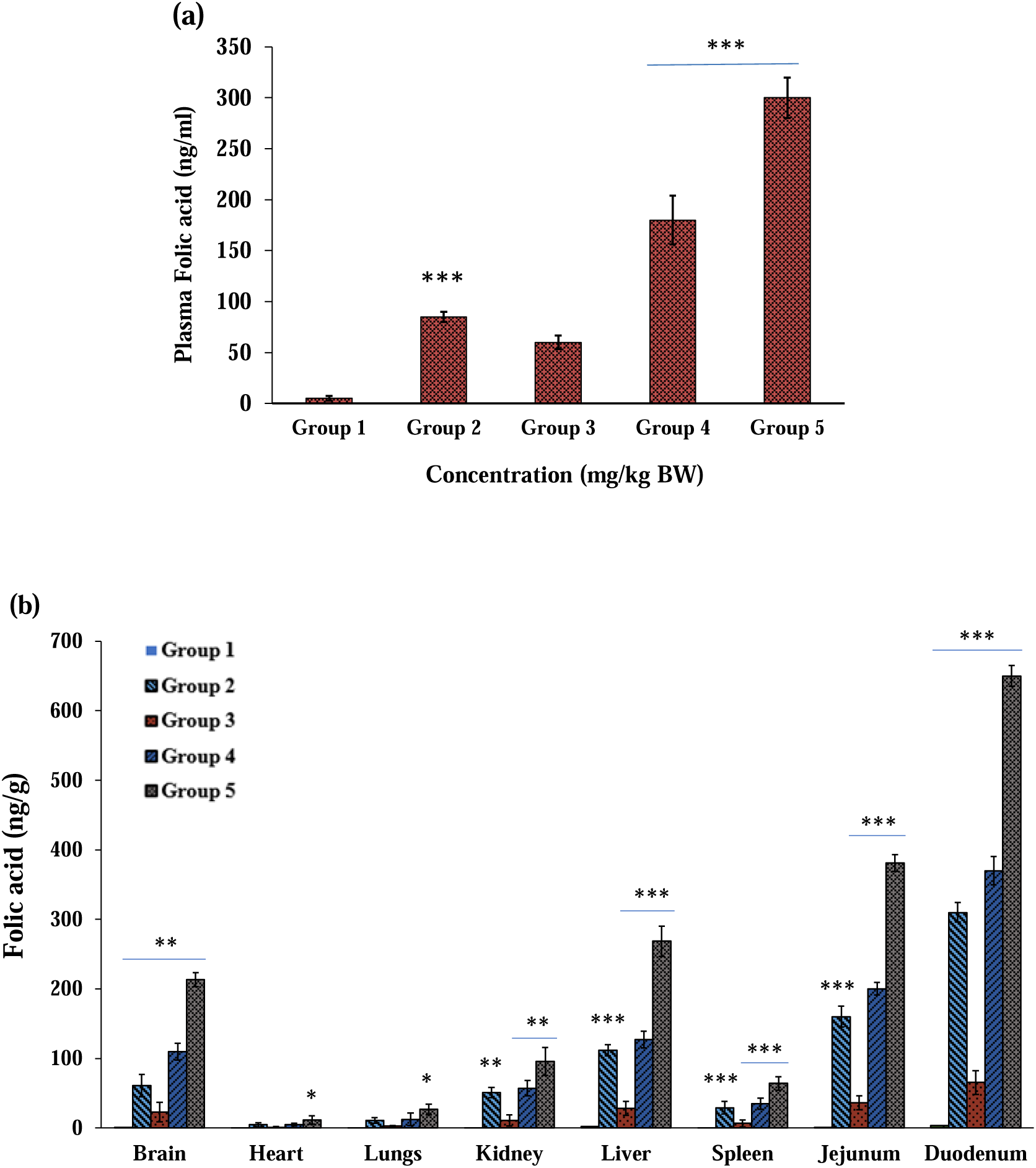
Bioavailability and tissue distribution of free FA and FA in FA-Chi-NPs fed rats; (a) Plasma FA levels after sub-acute doses (b) Tissue distribution profile of FA from FA-Chi-NPs after sub-acute doses. Group 1-control group (physiological saline), Group 2- (free FA - 4 mg/kg body weight), Group 3 (FA-Chi-NPs - 0.4 mg/kg body weight), Group 4 (FA-Chi- NPs - 2 mg/kg body weight), Group 5 (FA-Chi-NPs - 4 mg/kg body weight). Values are mean ± SE (n = 6). Values not showing a similar superscript are significantly different among the groups (One way ANOVA, *p < 0.05, ** p < 0.01, *** p < 0.001).

## 4. DISCUSSION

The potential for functional foods and nutraceuticals to contribute to healthy lifestyles has generated great interest in recent years. Their capacity to increase daily nutrient intake and prevent deficiencies may help to minimize the risk of various diseases (Annunziata et al., 2011). To develop functional foods, bioactive compounds such as antioxidants, minerals, and vitamins can be incorporated into food matrices (de Boer et al., 2016). However, in industrial contexts, where processing and storage can influence solubility, stability, and bioavailability, incorporating these compounds and maintaining their biological properties remains difficult. To overcome these constraints, encapsulating agents might serve as carriers or delivery systems. The development of natural food-grade carriers, termed “biocarriers,” is a major source of innovation for modern food technology. Hence, the present study aimed to encapsulate FA using a biodegradable polymer, chitosan.

In this study, FA-Chi-NPs were prepared and determined for particle size, zeta potential (ZP), polydispersity index (PDI), and entrapment efficiency because these are the determining variables of encapsulant bioavailability. The average particle size, ZP, and PDI of FA-Chi-NPs were 191.7 ± 2 nm, + 52 ± 4 mV, while PDI was 0.18 ± 0.1 as evaluated by DLS analysis (Figure 1a), and the % entrapment efficiency was found to be 80%, which was much higher than those reported in earlier studies (do Evangelho et al., 2019; Pamunuwa et al., 2020). The size and surface charge of the particle plays a vital role in the stability, efficacy, and bioavailability of FA in the body. Based on these results, it may be concluded that chitosan is a suitable carrier for the encapsulation of FA.

Hemolysis represents the rupture of red blood cells (RBCs) and results in releasing haemoglobin and other internal contents into the surrounding fluid. If a significant number of RBCs disintegrate in the body, it might result in severe pathological conditions, including anaemia, renal failure, and jaundice (Dobrovolskaia et al., 2009). All biological materials entering the bloodstream get in contact with RBCs; therefore, assessing the hemolytic potential of the biomaterials is essential. Preliminary experiments on the interaction of the developed nanoparticles with human blood components are of utmost importance since they can be employed as drug delivery systems and administered orally/intravenously. Thus, the hemolytic potential of FA-Chi-NPs was evaluated for concentrations between 200 and 800 μg/mL using a spectrophotometric method. The results of the hemolysis assay are revealed in Figure 2. It was observed that as the concentration of nanoparticles increases, the hemolytic percentage increases slightly. A sample is termed hemolytic if the hemolytic percentage is superior to 5%. The graph clearly shows that the hemolytic percentage was less than 5% for all three tested concentrations (200µg/mL, 400µg/mL, and 800 µg/mL). Hence, our study agrees with earlier findings that display heamocompatibility of chitosan nanoparticles tested on RBCs (Jesus et al., 2020). Thus, the hemolysis study demonstrates that no hemolysis was induced by FA-Chi-NPs and is suitable for systemic administration into the bloodstream.

The WST-1 metabolic activity assay was used to assess the cytotoxic profile of FA-Chi-NPs over a wide range of concentrations, as shown in Figure 3a. The concentration range studied was shown to be non-cytotoxic. The results showed that the cytotoxicity caused by various concentrations is negligible. Thus, incubation of Caco-2 cells with nanoparticles evoked no cell membrane damage, and a regular cell cycle was maintained. Similar results were reported by Jesus et al. (2020), who investigated the toxicity of chitosan NPs on RAW 264.7 cell lines. As a result, this study confirms that FA-Chi-NPs are not toxic to Caco-2 cells and can be used as a safe drug delivery mechanism in the cellular system.

The prepared FA-Chi-NPs were tested for ROS generation in Caco-2 cells. H_2_-DCFDA staining of cells showed a negligible increase in fluorescence intensity when treated with both free FA and FA-Chi-NPs. The measured concentrations of 200, 400, and 800 µg/mL did not reveal a dose-response in the formation of ROS, as shown in Figure 3c. On the contrary, positive control cells treated with H_2_O_2_ considerably increased ROS generation as compared to the control. Hence, these findings indicate that NPs are biocompatible and do not cause cell damage. In contrast, the concentration of 100 µg/mL and 400 µg/mL of free FA generated ROS slightly higher than control under the non-cytotoxic range. Earlier studies on the influence of chitosan nanoparticles on cellular ROS generation displayed conflicting results. Another study reported that chitosan NPs had a suppressive activity (Bor et al., 2016), whereas some investigations suggested no impact (Jesus et al., 2020; Omar Zaki et al., 2015), and several other research findings reported a stimulatory effect (Hu et al., 2011; Jiao et al., 2018; Sarangapani et al., 2018) on basal ROS cellular production. Further, the production of ROS results in the generation of lipid radicles, which can generate end-products such as MDA/DNA reactive aldehydes, which bind to DNA to form mutagenic adducts, thereby damaging membrane lipids (Venuprasad et al., 2013). Our results showed a non-significant increase in levels of MDA when cells were treated with different concentrations of free FA and FA-Chi-NPs, and the results are summarized in Figure 3b. Based on our findings, we concluded that there were no significant variations in the cytotoxic profile of chitosan NPs in Caco-2 cells which is in agreement with previous reports (Jesus et al., 2020).

Despite the numerous benefits of nano-formulation, the safety of the nanocarrier system and the type of interaction between the carrier and the bioactive molecule remain unknown (Lü et al., 2009). As a result, validating the toxicity of the nanocarrier system *in vivo* prior to application is necessary. As a result, the current study investigated the acute and sub-acute oral toxicity of FA-Chi-NPs in Wistar rats. After single oral administration of FA-Chi-NPs with variable doses (0.4 to 4 mg/kg body weight) in acute toxicity study, no adverse impact on general morphological, macro and microscopic structure of organs and tissues, haematological, plasma, and urine biochemical markers. Furthermore, there was no mortality or morbidity and no signs of toxicity, indicating that the LD_50_ value of FA-Chi-NPs in rats was greater than 4 mg/kg body weight, which is ten times higher than the daily recommended level of FA (0.4 mg) for a healthy individual. Similarly, Ravi and Baskaran (2015) also reported that fucoxanthin-loaded chitosan nanogels did not cause any clinical signs of toxicity or mortality. All factors were found to be within the normal laboratory reference range for rats, with no variations among the experimental groups.

Both acute and sub-acute oral toxicity investigations revealed no abnormal behaviour, growth, movement, morphology, feed, and water intake in comparison with control rats. The output of urine and feces was normal, with no signs of turbidity or diarrhea, indicating that the organs’ metabolic processes were unaffected. Further, the absolute and relative organ weights did not change significantly after receiving variable doses of FA-Chi-NPs. Additionally, the haematological parameters of rats showed no alterations, and the values reported were within the normal laboratory reference limits, displaying the safety of FA-Chi-NPs. Hence, the current finding indicates that FA-Chi-NPs had no toxicological impact. In mice, (Ranganathan et al., 2016) found that PLGA-PL NPs (1 and 10 mg/kg body weight) had no toxic effects. They believed that since lutein-poly- (lactic-co-glycolic acid) is a synthetic polymer, it could elicit physiological alterations after long-term feeding. Therefore, in the present study, we used natural biodegradable polymer instead of synthetic polymers Ravi and Baskaran (2015) reported no evident adverse effect of chitosan nanogels with fucoxanthin indicating that chitosan could be a suitable polymer matrix for drug delivery. Additionally, Pokharkar et al. (2009) also illustrate that chitosan gold nanoparticles exhibited no treatment-related toxicity in rats after oral administration, suggesting that it could be exploited for promising therapeutic applications. Ansari et al. (2019), Abdelazeim et al. (2020), and Shelat et al. (2018) reported that metallic NPs cause toxicity in rats because smaller particles have higher toxicity than larger particles. Since no adverse effects of FA-Chi-NPs were detected in this investigation, they were considered safe despite their smaller particle size. Thus, the FA-Chi-NPs developed in this study have the potential to be a unique FA delivery system.

Results from the current study demonstrate that liver enzymes such as AST ALP or ALT levels were not changed in all the tested doses (p < 0.05) in FA-Chi-NPs fed group. Moreover, the alterations (12%) in the AST activity in the chitosan-fucoxanthin nanogels treatment is observed by Ravi and Baskaran (Ravi et al., 2015) but were not validated by histology of the liver or specific liver functional indicators in plasma, such as ALP and bilirubin. However, the alterations were within the normal physiological range of AST and thus not related to nanogels. There was no significant change in renal functional markers such as uric acid, urea, or creatinine levels, indicating no variations in the kidney function of male and female rats. The findings of plasma chemistry examination from treatment groups show that administration of the FA-Chi-NPs at doses up to 4 mg/kg/day to rats for 28 days had no toxicologically significant harmful effects.

Histopathological examination of vital organs demonstrates unequivocally that FA-Chi-NPs had no toxicological impact on cellular structure. As a result, the no-observed adverse effect limit (NOAEL) of FA-Chi-NPs (FA equivalent) investigated in the 28 days of study was higher than or equal to 4 mg/kg BODY WEIGHT, the maximum dose tested in both male and female rats. However, this is the first report to confirm the safety assessment of FA loaded in a natural polymeric nanocarrier.

Oxidative stress has been identified as a potential mechanism of nanomaterials toxicity (Ahamed et al., 2011). This study believed that oxidative stress is a critical factor in nanoparticle-induced toxicity in experimental rats. It might interrupt the antioxidant defence mechanisms of the liver by altering antioxidant-related enzyme activities (Cao et al., 2016). Free radicals accumulate when the body’s antioxidant activity no longer protects the cell from oxidative stress, leading to adverse effects, such as lipid peroxidation (LP) and protein oxidation (Zhou et al., 2015). One of the major contributors to cell membrane damage is LP, which is a process that promotes the formation of oxygen radical-related damages (Koc et al., 2003). LP represents the primary indicator of oxidative damage. Following the initiation of LP, a sequence of free radical-mediated reactions produces many toxic by-products, including MDA (Prabhakar et al., 2012). Thus, in this study, as shown in Figure 7a, the daily consumption of free FA and FA-Chi-NPs for 28 days had no effects on the MDA levels compared to the control group.

Nitric oxide (NO) levels and oxidative stress were also reported to play a major role in the mechanism of toxicity for a variety of NPs, leading to Kupfer cell activation and recruitment of inflammatory cells, which promotes oxidative damage via responsive transcription factor (Ashkenazi, 2002). The effect of NPs treatment on NO levels in tissues (liver, kidney, spleen) of experimental rats is summarized in Figures 7b and 8b. In our study, no alterations in NO level were observed in any of the treated groups compared to the control group.

Additionally, the exposure of rats to different concentrations of FA-Chi-NPs did not affect GSH levels (Figures 7c and 8c). This study is supported by the earlier findings where chitosan-loaded silver nanoparticles did not cause any changes in the levels of GSH (Hassanen et al., 2019). GSH protects cells from oxidative stress and serves as a major non-enzymatic aqueous-phase antioxidant and a necessary cofactor for antioxidant enzymes involved in cellular redox processes. The endogenous antioxidant mechanism, which includes SOD, CAT, TAC, GR, and GPx, plays a vital role in free radical and peroxide metabolism and is partly responsible for protecting cells from oxidative stress (Wang et al., 2006). In this study, we observed no significant increase in SOD, CAT, TAC, GR, and GPx activity (Figures 7d-7h and 8d-8h) in all treated groups compared to the control in acute and sub-acute toxicity studies.

Furthermore, only a few investigations were available on the safety aspects of natural polymer nanocarrier systems. Therefore, this is the first report exhibiting the toxicity of nanocarrier system for FA with bio-polymer, i.e., chitosan. As a result, it is crucial to investigate nanocarrier toxicity because it is one of the most important aspects to consider when assessing the potential of a novel drug delivery system. On the other hand, plasma levels of FA from NPs for sub-acute doses of FA-Chi-NPs (2, 4 mg/kg body weight) were significantly greater than those observed in a free FA dose (4 mg/kg body weight). Toragall et al. (2020) also described a similar profile of dose-dependent increase of lutein in plasma of rats fed with lutein-loaded chitosan-sodium alginate nanocapsules. The probable reason for the improved bioavailability of FA from Chi-NPs could be a slow and controlled release of FA from natural polymer chitosan. The biodistribution pattern of FA in rat organs was as follows: heart < lungs < spleen < kidney < brain < liver < jejunum < duodenum. Bernstein et al. (1970) demonstrated that the hydrolysis and absorption of FA polyglutamate require the presence of folate conjugase in the small intestine for converting polyglutamate into monoglutamates. Hence, FA absorption takes place primarily in the duodenum and jejunum. FA is not methylated after absorption in human and canine portal blood, but it may be methylated later in the liver after reduction. As a result, the present study observed significant folate deposition in the duodenum and jejunum tissues. Based on these results, the chitosan biopolymer used in the present work may have influenced the enhanced solubility, stability, and absorption of FA from Chi-NPs. To our knowledge, there is limited data available in the scientific literature on the bioavailability and tissue distribution of FA after sub-acute doses of FA-Chi-NPs. Therefore, the current findings show that the chitosan nanocarrier could be a more efficient carrier for the delivery of FA for treating several disorders.

## 5. CONCLUSION

In this investigation, the FA-Chi-NPs were successfully fabricated by the ionic gelation method with 80% drug loading using TPP as a cross-linker. The NPs’ particle size and charge were assessed. The *in vitro* cytotoxicity assays show that the prepared NPs have no significant cytotoxic effect on Caco-2 cells and also proved its haemocompatibility. Toxicity studies of acute and sub-acute doses of FA-Chi-NPs exhibited no toxic effect in male and female Wistar rats. Sub-acute toxicity study displayed that the NOEL of FA-Chi-NPs was 4 mg/kg body weight, which was validated via haematology, plasma biochemical parameters, and histopathological investigations. Additionally, Chi-NPs were found to be safe since they did not show any alterations in the oxidative stress biomarkers. Furthermore, enhanced bioavailability and tissue distribution of FA were prominent from FA-Chi-NPs. However, this is the first study to demonstrate the safety and bioavailability of FA-Chi-NPs after acute and subacute doses. Therefore, chitosan might be employed as a model biopolymer for FA delivery and has been found to be a safe application in the pharma and food industries.

## ACKNOWLEDGEMENT

The authors would like to express their gratitude to the Director of the DRDO-Defence Food Research Laboratory for his unwavering support and encouragement throughout the research period.

